# Hairpin: a tool for predicting structural non-coding RNA genes *ad initio*

**DOI:** 10.1101/640862

**Authors:** Jakob Peder Pettersen

**Affiliations:** Department of Biotecnology and Food Science, NTNU - Norwegian University of Science and Technology, Sem Sælandsvei 6-8, 7491 Trondheim, Norway

**Keywords:** hairpin, ncRNA genes, gene predection, structural RNA, stem loops

## Abstract

**Background:** Structural RNA genes play important and various roles in gene expression and its regulation. Finding such RNA genes in a genome poses a challenge, which in most cases is solved by homology approaches. *Ab intio* methods for prediction exist, but are not that much explored.

**Results:** We introduce `hairpin` which identify potential structural RNA genes only based on the sequence. We use the algorithm to predict RNA genes in *Escherichia coli* K-12. When looking at very short regions of the genome, we do not get results differing very much from a random shuffling of the genome. However, at longer stretches it is a clear biological signal. It turns out that none of the regions predicted to code for RNA genes have such an annotation in literature.

**Conclusions:** Arbitrary DNA sequences seem to give rise to transcripts with secondary structures similar to real ncRNA. We therefore conclude that exclusively looking at secondary structure base-parings is in general a futile approach.

## Background

mRNA, the most known form of RNA, codes for proteins. The rest of the RNA is called non-coding. Nevertheless, non-coding RNA (ncRNA) plays important roles in gene expression and regulation thereof. The most known examples are tRNA and rRNA, both participating in proteins synthesis. Other examples include riboswitches in bacteria and miRNA in eukaryotes, which are examples of regulatory RNA. A region in the genome coding for ncRNA is called a RNA gene. Due to the their importance and various functions, finding RNA genes and their functions might help us to get a better understanding of the organism in question. Unlike protein-coding genes, RNA genes cannot be found by searching for open reading frames, making them harder to predict [1, 2].

Many ncRNA molecules have structures and motifs required for their function. In this respect, we mainly focus on the secondary RNA structure, this is the two-dimentional structure formed by base-pairings between nucleotides in the same molecule. A RNA gene where the secondary structure is related to its function, is called structured *sensu* Meyer [2]. Three different base-pairings occur in RNA molecules. In addition to the Watson-Crick pairs {A,U} and {G,C}, we also have the {G,U} base pair [2]. One of the most common motifs being observed in structural RNA molecules is the stem-loop, where a stem consisting of two anti-parallel parts of the molecule base-pair, where the two parts of the stem is separated by a loop comprising a few nucleotides [3].

Most approaches for RNA gene prediction are based on homology [2] where similarities in sequence or structure to known ncRNAs are used. Sometimes however, we may want to discover new RNA genes where information from homology is lacking. Such *ab initio* methods are rarer and considered more challenging [2]. There exist some specific methods based on nucleotide composition in AT-rich hyperthermophiles [4] and methods dedicated to find specific kinds of RNA genes [5]. A more general prediction method introduced by [6] uses the fact that base pairings lower the *minimum free energy* involved in RNA folding. The current article’s idea of using hairpins is not new either, [7] discusses this idea, but the level of detail is insufficient to get a clear overview of the procedure.

## Implementation

We introduce `hairpin`, a command-line application for Python 3.5.0 or later, using Biopython [8] and NumPy [9]. The main idea of `hairpin` is to identify potential hairpins in the input and group them together. First, the entire nucleotide sequence is transcribed into a continuous RNA strand, using the input as the coding strand. The main idea is that structural RNA genes contain more and longer hairpins than expected by chance, distinguishing them from other parts of the genome. The algorithm thus has to consider the following:

- How do we find the hairpins?
- What do we mean by “random chance”?
- Once the hairpins are found, how do we use them to predict which parts of the genome contain RNA genes?

Different from [10], we do not seek to filter out RNA being regulatory parts in protein coding genes, such as attenuation sequences.

Note that the pseudocode written in this article does *not* directly reflect the actual implementation as it contains many optimizations which would be difficult to understand. Instead, the pseudocode aims to produce the same results.

### Finding the hairpins

When searching for hairpins, we look for base-pairings involving consecutive residues. Taking figure 1 as an example; we observe that residues 1-5 (from the lower left) base-pair with residues 11-14, forming a stem. We note the following: Residue 1 can base-pair with residue 14, being 13 nucleotides apart, residue 2 can base-pair with residue 12, being 11 nucleotides apart, residue 3 can base-pair with residue 11, being 9 nucleotides apart and so on. `hairpin` utilizes this observation such that consecutive nucleotides having potential base-paring partners at a decreasing distance (decrease with two for each nucleotide), is considered a motif. We hence include the parameter *l*_*min*_ telling the minimum length of such a motif in order to report it. Setting *l*_*min*_ to a higher value makes it more difficult for motifs to arise by pure chance.

**Figure 1:**
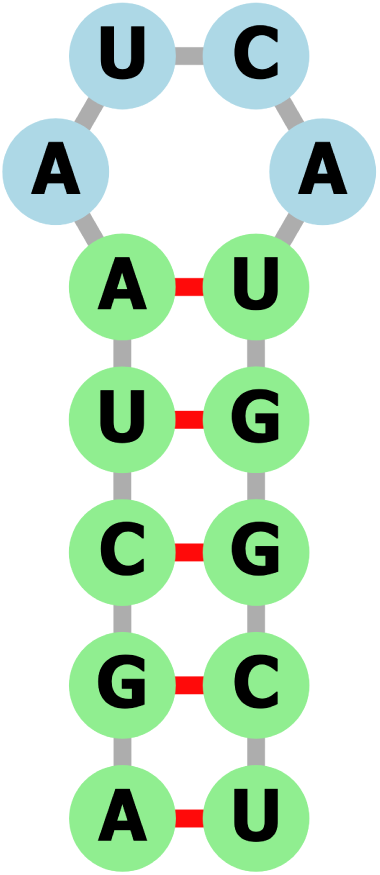
Sample RNA stem-loop structure, generated with `forna` [11]

The pseudocode for this procedure is shown in algorithm 1.

### Estimating the structuredness of a RNA region

Telling whether a region of the RNA transcript is likely to represent structural RNA, requires some way to quantify the “structuredness” of the region. This will be based on the motifs occurring in the region. Instead of using a sophisticated algorithm to predict how the strand will fold as done by [6], we naively look at the number of nucleotides in the region taking part in one or more motifs, hereafter called the base-pairing count. We do not know whether there exists that many nucleotides being able to base-pair in the real RNA structure as the motifs may overlap, but the same nucleotide is never counted twice. However, this approach is meant as a heuristic being easy to compute. The pseudocode is listed in algorithm 2.

### Computation of randomizations

Randomized RNA segments are built by sampling nucleotides with replacement from the original transcript. We designate parameters *r*_*min*_ and *r*_*max*_ being respectively the minimum and maximum length of the RNA regions to probe. For each length *r* in the closed interval [*r*_*min*_, *r*_*max*_], we take the *r* first nucleotides and find the hairpins and base-pairing count. The procedure is repeated *n*_*rand*_ times with newly generated RNA segments. Finally, for each length *r*, the mean base-paring count and its standard deviation is computed. See algorithm 3 for pseudocode.

### Comparing actual motifs found to the randomized results

We screen trough the motifs for finding regions of appropriate size. If we find a such a region, we report the base-pairing count. This count is compared to what to expect by random using the assumption that this quantity is normally distributed under the null hypothesis. From this, a *p*-value is computed. This *p*-value is corrected by the length of the original sequence by the following logic:

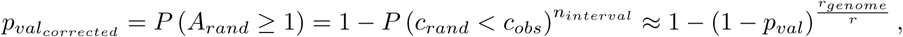

where:

- *A*_*rand*_ is the number of regions of length *r* in the genome having an equal or higher base-paring count than *c*_*obs*_ under the null hypothesis
- *c*_*rand*_ is the base-paring count in a interval of length *r* under the null hypothesis
- *c*_*obs*_ is the observed base-paring count
- *n*_*interval*_ is the number of disjoint regions in the genome of length *r*
- *r* is the length of the region
- *r*_*genome*_ is the length of the genome

Thereafter, we go through the intervals, pick the most significant ones and discard the intervals overlapping with it. Furthermore, we account for multiple testing by correcting with the Benjamini–Yekutieli procedure [12], computing *q*-values. We keep all regions with *q*-value less than *q*_*crit*_. The procedure is listed in algorithm 4. The results are finally reported as:

- A `.bed` track file to be uploaded in a genome browser. It includes the start and end position of significant regions being found, in addition to their *q*-values.
- A text file displaying the results for the randomized results with the mean base-pairing count for each length of the regions, together with its standard deviation.
- A plot showing the means and standard deviation of the base-pairing counts of the randomized results together with the lengths and base-pairing counts of the significant regions found.

## Results

Using the parameters *d*_*min*_ = 4, *d*_*max*_ = 25, *q*_*crit*_ = 0.01, *l*_*min*_ = 8 and *n*_*rand*_ = 2000, the algorithm was run on the genome of *Eschericia coli* K12 [13], with four variations:

- *r*_*min*_ = 25 and *r*_*max*_ = 200 on the original genome
- *r*_*min*_ = 25 and *r*_*max*_ = 200 on a random reshuffling of the genome
- *r*_*min*_ = 200 and *r*_*max*_ = 1000 on the original genome
- *r*_*min*_ = 200 and *r*_*max*_ = 1000 on a random reshuffling of the genome

Running all the jobs in parallel on a machine with 48 2.4 GHz cores took 8501 seconds. Plots displaying the results are shown in figure 2.

**Figure 2:**
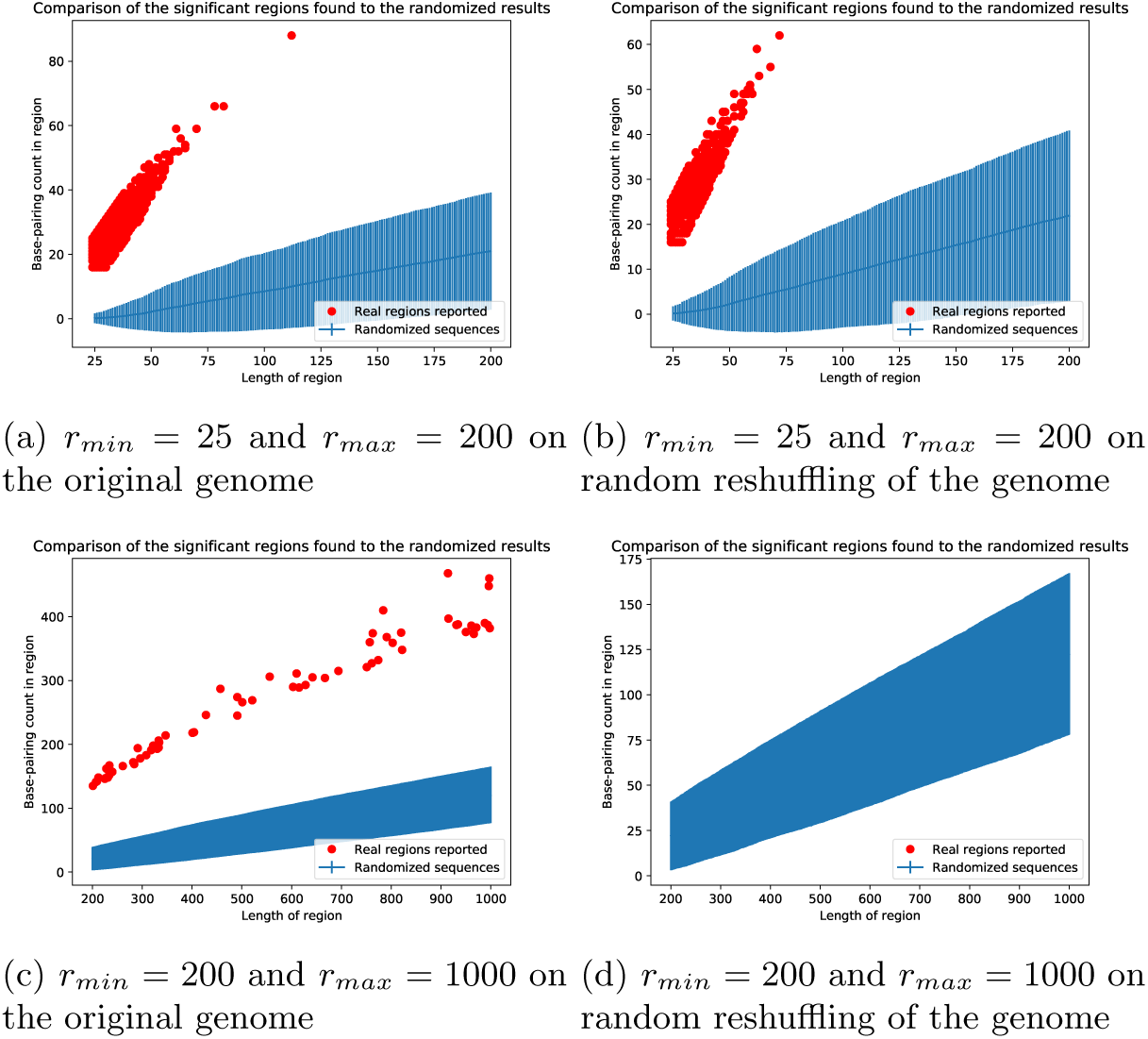
Each significant region found is marked with a red dot, telling its length and its base-paring count. The results for the randomized sequences are shown in blue where the blue line is the mean base-pairing count and the error bars correspond to the standard error. For *r*_*min*_ = 200 and *r*_*max*_ = 1000, the error bars are too dense to be seen.

The number of significant regions found is shown in table 1. For *r*_*min*_ = 200, *r*_*max*_ = 1000 on the original genome, the genome track file was uploaded to the UCSC Microbial Genome Browser [14] and the most significant regions were looked up manually. The results thereof are shown in table 2. Furthermore, the RNA corresponding to the first region listed in table 2 was visualized with `forna` [11] being part of the ViennaRNA Web Services, along with a transcript being of the same length being built from a reshuffling of the nucleotides of the genome. This is shown in figure 3.

**Table 1:** Number of significant regions found for each of the runs

| Genome | $r_{min}$ | $r_{max}$ | Number of significant regions found |
| --- | --- | --- | --- |
| Original | 25 | 200 | 12791 |
| Shuffled | 25 | 200 | 8400 |
| Original | 200 | 1000 | 67 |
| Shuffled | 200 | 1000 | 0 |

**Table 2:**
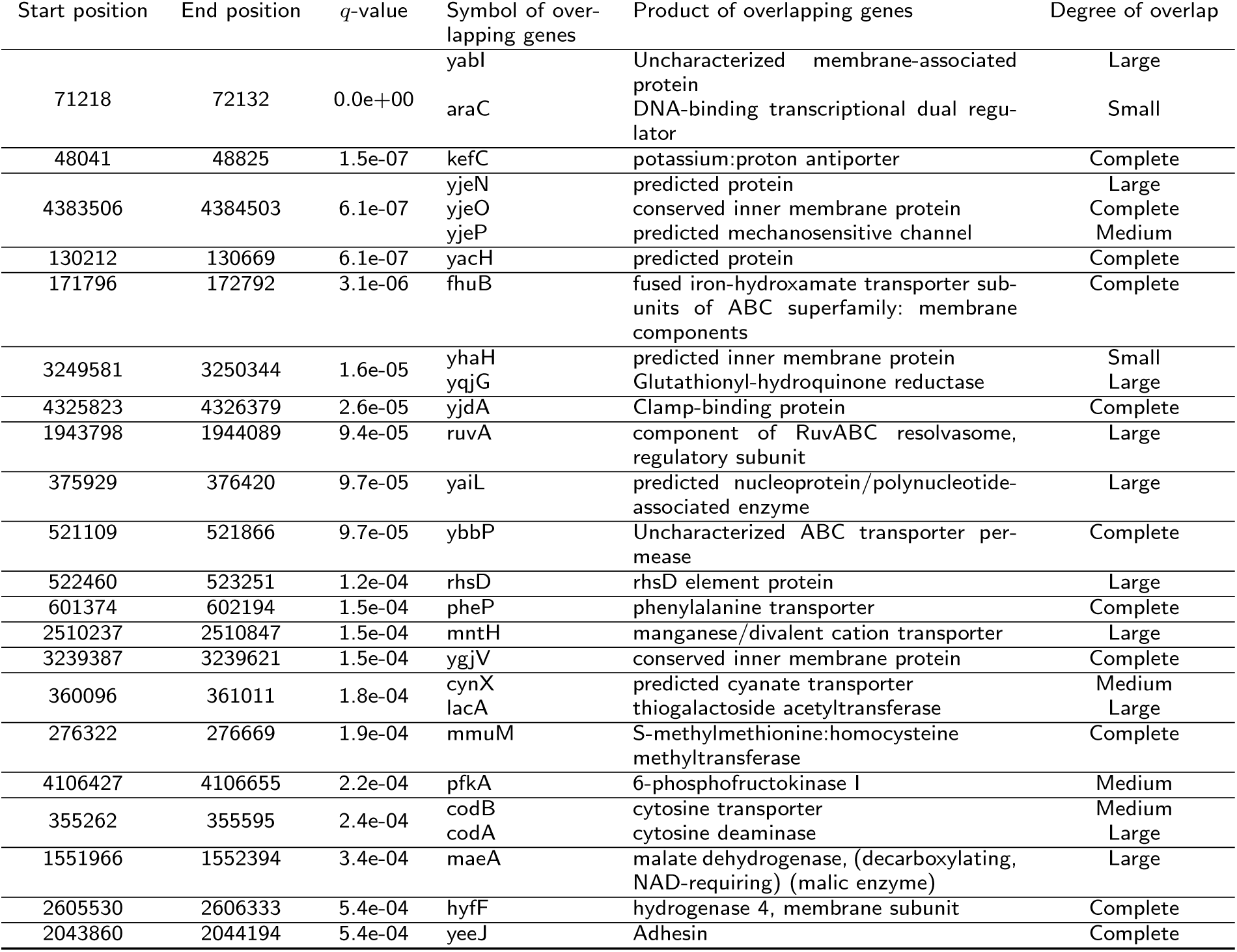
Position, *q*-value and overlapping genes for the most significant regions found with *r*_*min*_ = 200, *r*_*max*_ = 1000 on the original genome

**Figure 3:**
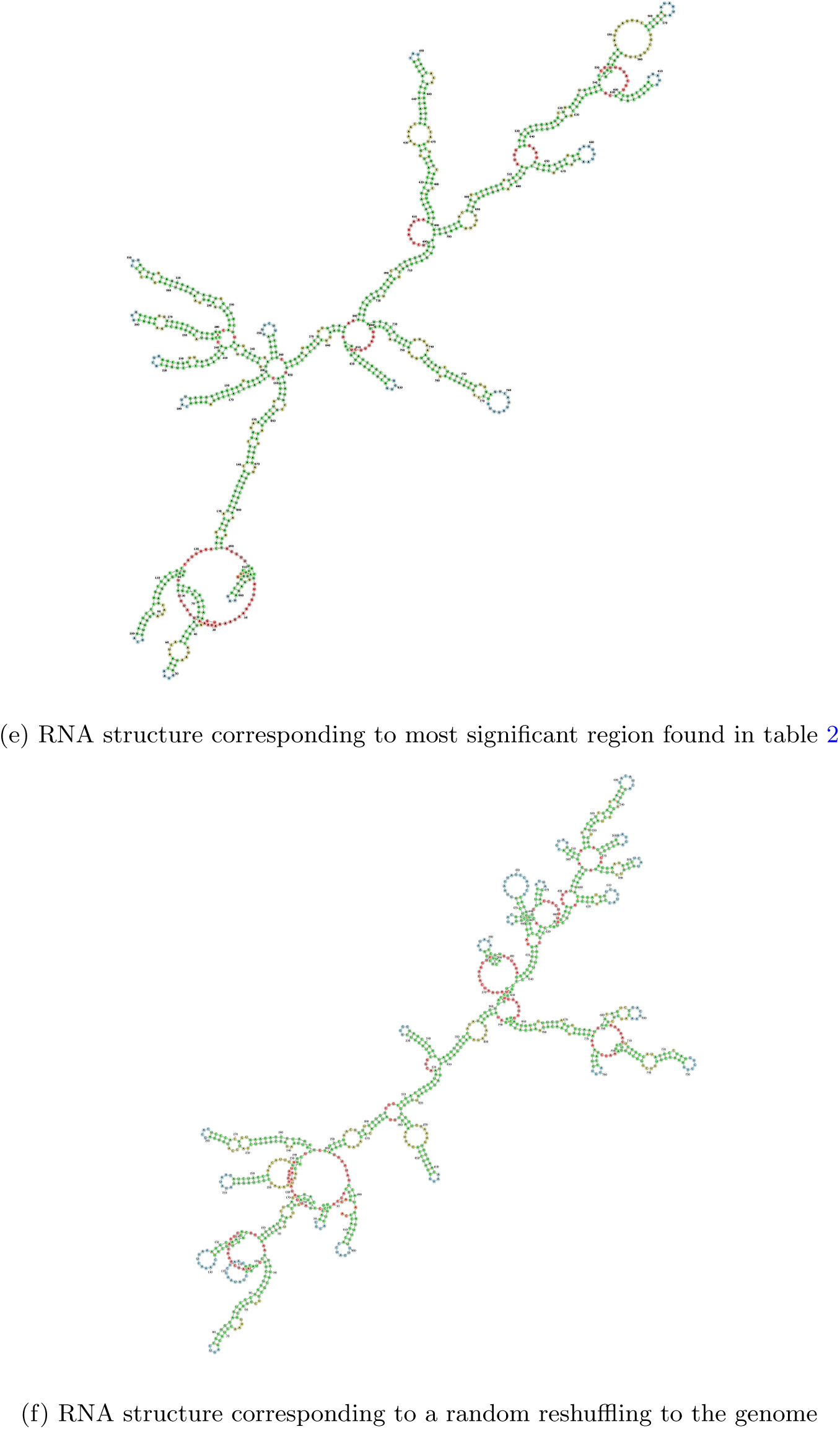
RNA structure generated with `forna`

## Discussion

For *r*_*min*_ = 25 and *r*_*max*_ = 200, significant regions were found also when randomly shuffling the genome. Hence, the results for these parameter value should not be trusted and suggest that the method of correcting for the *p*-values is not appropriate. However, when increasing the size of the regions of search (*r*_*min*_ = 200 and *r*_*max*_ = 1000), we see positive results for the real genome, but not the shuffled one. This indicates that our algorithm detects a real biological signal. However, when inspecting table 2, we discover that our findings are not as expected. None of the significant regions shown seem to correspond to the Genebank ncRNA genes annotated in the genome browser, nor do the results seem to agree with [10]. The genes overlapping with significant regions are of various types, but seem to be enriched in membrane transporter proteins.

Surprisingly, figure 3 shows no big difference in the folding pattern of the most significant region in table 2 and a random transcript. This leads us to think that RNA strands being completely random also may attain secondary structures resembling those of real ncRNAs. Hence, we it seems likely that structural features does not distinguish ncRNA genes from other genomic elements, agreeing with [15].

According to [4] an integrated approach taking several kinds of information into account, is probably a more fruitful approach. For instance [16] uses thermodynamic stability (based on secondary structure) in addition to evolutionary multiple alignments and obtains results which are very good. Hence, we might think the same about `hairpin`: Even though the procedure by itself is insufficient to predict ncRNA genes, it can probably be used in combination with other methods in order to obtain more accurate and robust results.

The bias resulting in the difference between figure 2c and figure 2d still needs an explanation. It could due to features such as repeats or a special composition of nucleotides in the real biological sequences.

## Conclusion

Hoping to predict ncRNA genes *ab inito* using the secondary structure only, turned to be too optimistic. Even though the secondary structure is essential for the function of ncRNA, their structures seem to drown in random noise. There is hope however, that when used in combination with other kinds of information, `hairpin` can be useful to predict ncRNA genes.

## Competing interests

The author declare that they have no competing interests.

## Author’s contributions

Not applicable

## Availability of data and materials

The genome used in the analysis is the complete genome of *Escherichia coli* strain K-12 substrain MG1655, available as NC 000913.3 from RefSeq. At http://microbes.ucsc.edu, the same assembly is available in the genome browser. The software in addition to the analyses being run in this experiment are found on https://github.com/yaccos/hairpin.

### Acknowledgements

Thanks to Finn Drabløs for providing positive feedback to this article.

## Pseudocode

### Algorithm 1 Method for finding hairpins in a RNA transcript

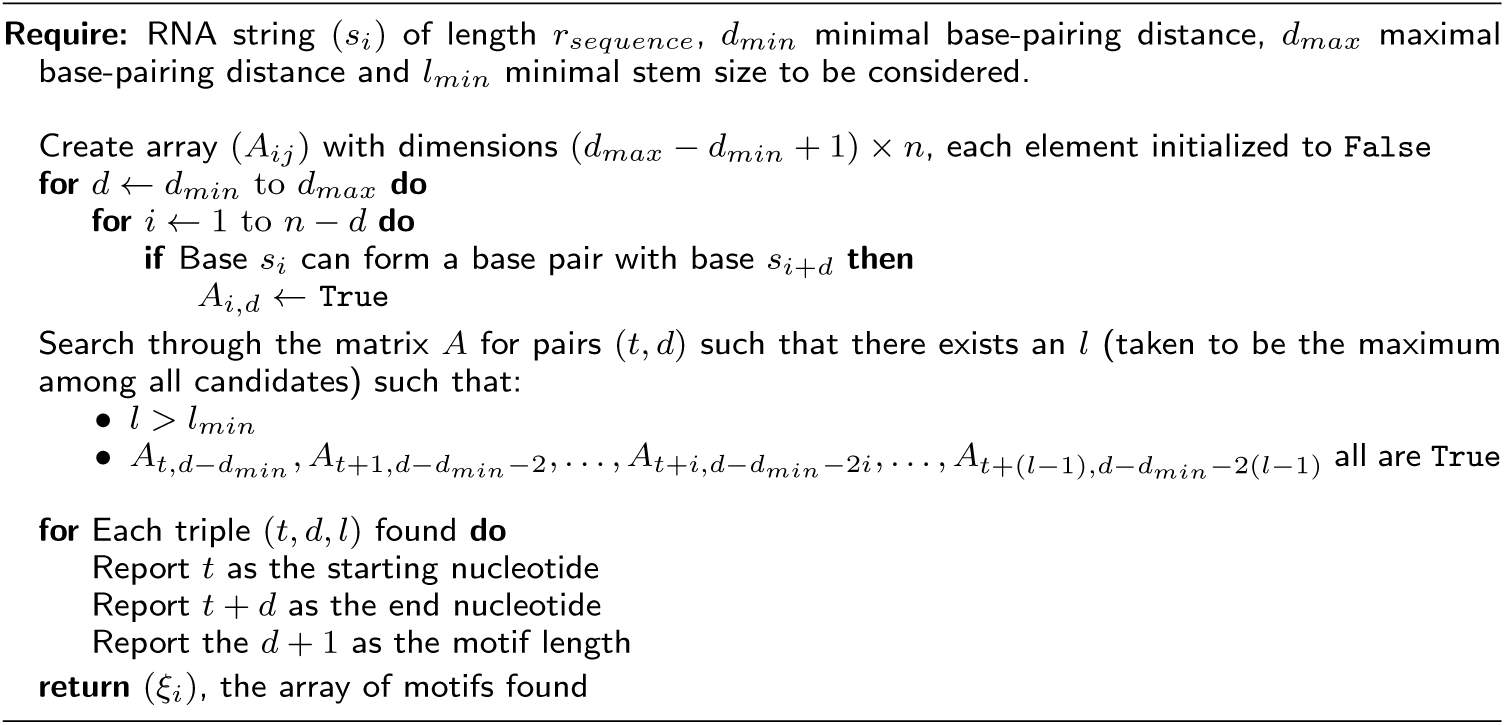

### Algorithm 2 Method for computing base-pairing count

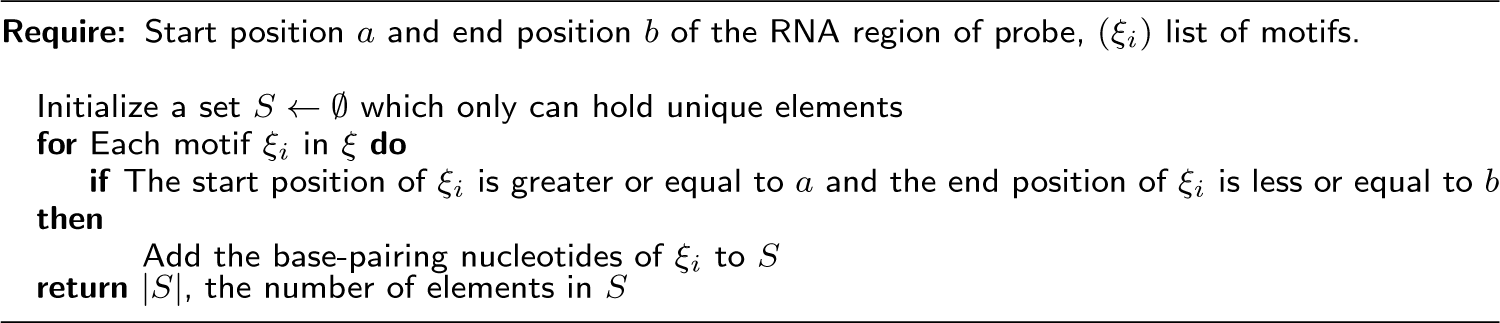

### Algorithm 3 Computing randomized RNA strands

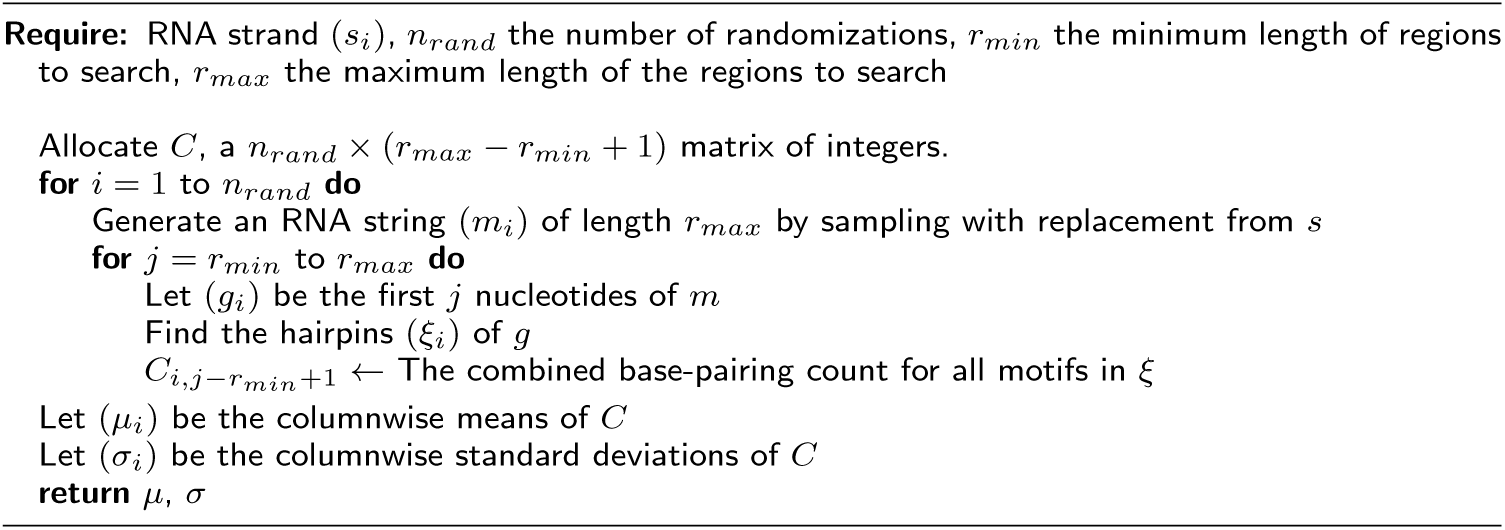

### Algorithm 4 Finding putative ncRNA coding regions

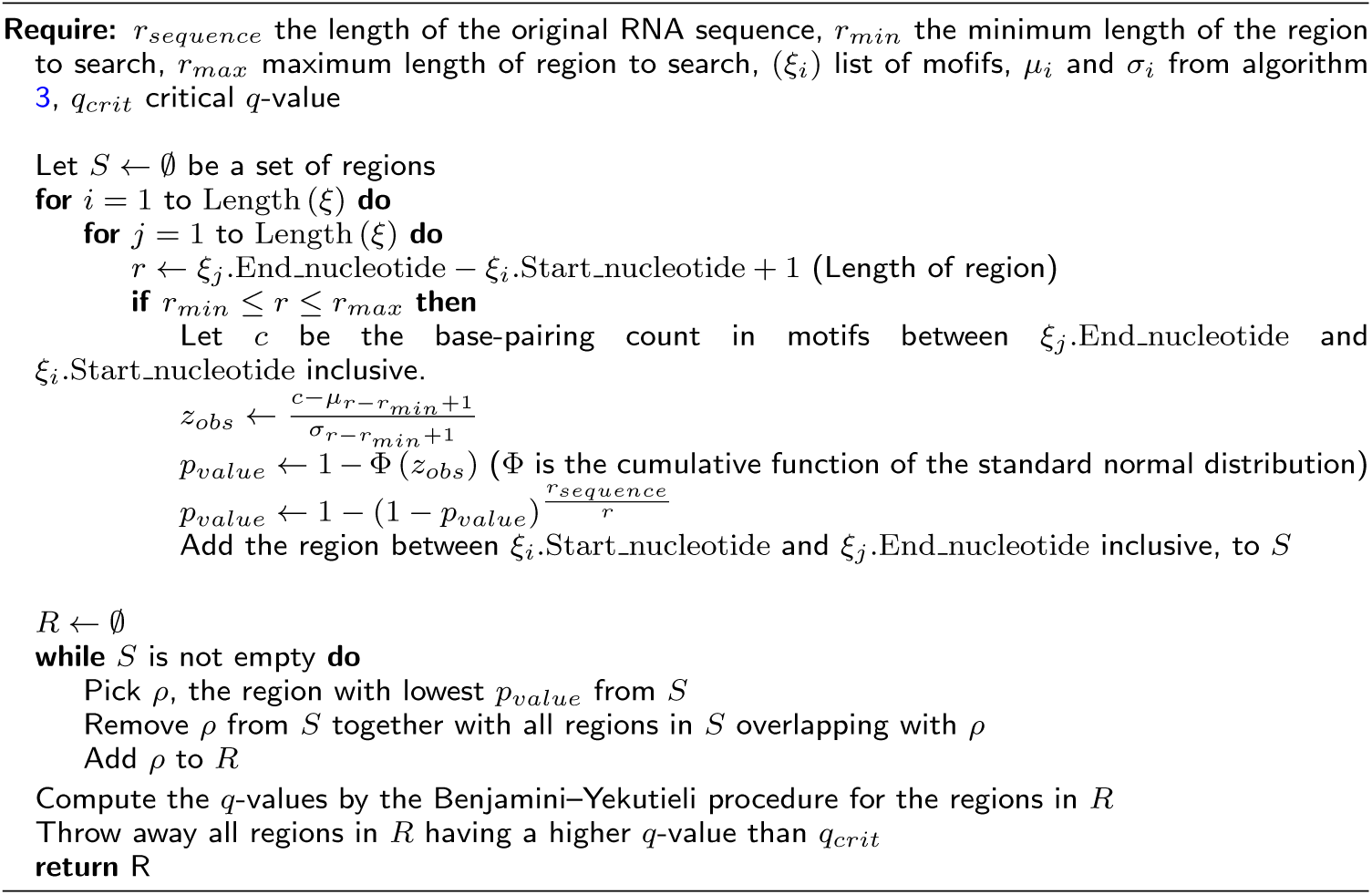

## Notes

https://github.com/yaccos/hairpin

